# Single-cell Characterization of Acute Myeloid Leukemia and its Microenvironment Following PD-1 Blockade Based Therapy

**DOI:** 10.1101/2020.09.03.278309

**Authors:** Hussein A. Abbas, Dapeng Hao, Katarzyna Tomczak, Praveen Barrodia, Jin Seon Im, Patrick K. Reville, Zoe Alaniz, Wei Wang, Ruiping Wang, Feng Wang, Gheath Al-Atrash, Koichi Takahashi, Jing Ning, Maomao Ding, Jairo T. Mathews, Latasha Little, Jianhua Zhang, Sreyashi Basu, Marina Konopleva, Guillermo Garcia-Manero, Michael R. Green, Padmanee Sharma, James P. Allison, Steven M. Kornblau, Kunal Rai, Linghua Wang, Naval Daver, Andrew Futreal

**Author notes:** These authors contributed equally: Hussein A. Abbas, Dapeng Hao, Katarzyna Tomczak. Co-corresponding authors, **Kunal Rai, PhD**, Associate Professor, Department of Genomic Medicine, M D Anderson Cancer Center, Houston, TX 77030, Tele, **Linghua Wang, MD, PhD**, Assistant Professor, Department of Genomic Medicine, M D Anderson Cancer Center, Houston, TX 77030, Tele, **Naval Daver, PhD**, Associate Professor, Department of Leukemia, M D Anderson Cancer Center, Houston, TX 77030, Tele, **Andrew Futreal, PhD**, Professor and Chair, Department of Genomic Medicine, M D Anderson Cancer Center, Houston, TX 77030, Tele.

## Abstract

Acute myeloid leukemia (AML) and effector cells of immune checkpoint blockade (ICB) therapy co-reside in a complex bone marrow (BM) milieu. The interplay of tumor intrinsic and microenvironment (TME) mechanisms that influences the response to ICB-based therapies in AML have not been elucidated. Here we report our analyses of single cell RNA profiling of more than 127,000 BM cells from healthy donors and relapsed/refractory (R/R) AML patients at pre/post treatment with azacitidine/nivolumab, paired with single cell T cell receptor (TCR) repertoire profiles, to uncover factors impacting response and resistance. Loss of chromosome 7/7q conferred an immunosuppressive TME and was associated with resistance to ICB-based therapy in R/R AML. Our trajectory analysis revealed a continuum of CD8+ T cell phenotypes, characterized by differential expression of granzyme B (GZMB) and GZMK. GZMK expression defined a BM residing memory CD8+ T cell subset with stem-like properties likely an intermediary between naïve and cytotoxic lymphocytes. Responses to ICB-based therapy were primarily driven by novel and expanded T cell clonotypes. Our findings support an adaptable T cell plasticity in response to PD-1 blockade in AML. Disentangling AML cells from their complex, immune-rich microenvironment revealed characteristics that shaped resistance to ICB-based therapy and could inform strategies to target AML vulnerabilities.

**Significance:** Determining the cellular and molecular underpinnings of response and resistance to PD-1 blockade based therapy in AML can guide immune-based therapeutic strategies. Our results reveal AML intrinsic characteristics (chromosome 7/7q status and oxidative stressors) and tumor microenvironment to modulate responses to checkpoint blockers. CD8 cells exist in the bone marrow in a continuum with GZMK expression defining a memory, stem-like T cell population that could play a role in response to therapy.

## Introduction

The BM constitutes the TME of AML, harboring complex cellular components in different states of differentiation which may impact response to treatment(1,2). About 25-30% of patients with AML are refractory to frontline therapy and among those who respond another 50-60% will relapse(3). Patients who are refractory or relapse within 6 months post frontline induction therapy have poor outcomes with low remission rates and median survival of 4-6 months with cytarabine or hypomethylating agent-based salvage therapies(4–6). Allogeneic hematopoietic stem cell transplantation (ASCT) remains the only curative option for patients with R/R AML, largely achieved via the induction of allo-reactive T cells against residual leukemic cells via the graft versus leukemia effect(7). The success of ASCT in curing AML provides strong evidence that the immune system can be harnessed to eradicate AML. ASCT may not be available to many AML patients due to older age, patient comorbidities, donor availability, cost, and access to centers of expertise.

Building on the success of ASCT in AML and bispecific antibodies in B-ALL, off-the-shelf immunotherapies such as immune checkpoint blockade (ICB) based therapies and bispecific antibodies are being developed in AML(8,9). Cytotoxic T-lymphocyte-associated protein 4 (CTLA4) blockade induced encouraging and durable responses in post-ASCT relapsed AML(10). In a phase 2 trial in R/R AML, the combination of the anti-programmed cell death protein 1 (PD-1) agent nivolumab in combination with the hypomethylating agent azacitidine demonstrated improved response rates and median overall survival compared with a contemporary cohort of similar patients treated on azacitidine-based clinical trials(11). Through these and other ongoing clinical trials a marked variability in the efficacy of ICB based therapies in patients with AML has been noted(10,11). Furthermore, both the median duration of response and the proportion of long term responders was lower in R/R AML compared to what has been reported for ICB treatments in many solid cancers(10–15). These observations suggest that thus far undefined intrinsic and TME resistance mechanisms may impede ICB efficacy in AML.

Single cell profiling of solid tumors and their associated TME have identified unique cellular subsets and pathways associated with response to ICB. For instance, melanoma resident TCF7+ CD8+ cells, abundance of dysfunctional CD8+ T cells, and accumulation of exhausted T cells in melanoma correlated with improved responses to ICB-based therapies(16–19). Furthermore, both interferon gamma (IFNγ) and oxidative phosphorylation pathways affected response to ICB(20–22). Single cell T cell receptor (scTCR) repertoire analysis revealed the expansion of novel clones in basal cell and squamous cell carcinoma in response to PD-1 blockade(23). Whether these mechanisms also exist in the complex BM milieu of AML patients and modulate ICB based therapy responses remain largely unknown.

We conducted longitudinal single cell RNA (scRNA) profiling of 113,394 BM cells from 8 patients with R/R AML treated with azacitidine and nivolumab therapy to understand the tumor intrinsic and TME factors involved in response, resistance, and relapse to this therapy. We also investigated the scTCR repertoires of treated patients to characterize the lineage and functional changes in T cells that were associated with response to ICB-based therapy. These results afford a deep characterization of the BM TME in AML in the context of ICB-based therapy, providing important insights upon which to further improve this modality for leukemia patients.

## RESULTS

### Patient cohort and characteristics

Eight pretreatment and 14 posttreatment BM aspirates from 8 patients with R/R AML (median age 73 years; range 64-88 years) treated with azacitidine and nivolumab (hereafter referred to as ICB-based therapy) on NCT02397720 were profiled with scRNA sequencing (scRNAseq) and paired with scTCR sequencing (Fig. 1a). Patients with R/R AML, of any age, with adequate organ function and ECOG performance status 0-2 were eligible for the trial (complete clinical protocol as Supplemental file). Three patients (3/8) had complete/partial response including 2 CR and 1 PR (hereafter referred to as responders), 2/8 patients had stable disease (SD) and 3/8 patients had no response (NR) to ICB-based treatment, per European Leukemia Net (ELN) 2017 response criteria(24). Patient and response characteristics are shown in Supplementary Figure 1a and Supplementary Table 1. Seven of 8 patients had previously received and progressed on hypomethylating agents (azacitidine or decitabine). Six of 8 patients had at least one cytogenetic abnormality prior to treatment initiation, including 3/3 NR patients who had chromosome 7/7q deletion. Targeted sequencing in at least 1 timepoint per patient (total evaluated 17/22 timepoints) revealed mutations in *ASXL1* (4/8 patients), *TET2* (3/8 patients), *SRSF2* (3/8 patients) and *FLT3* (2/8 patients) (Supplementary Figure 1b).

**Figure 1.**
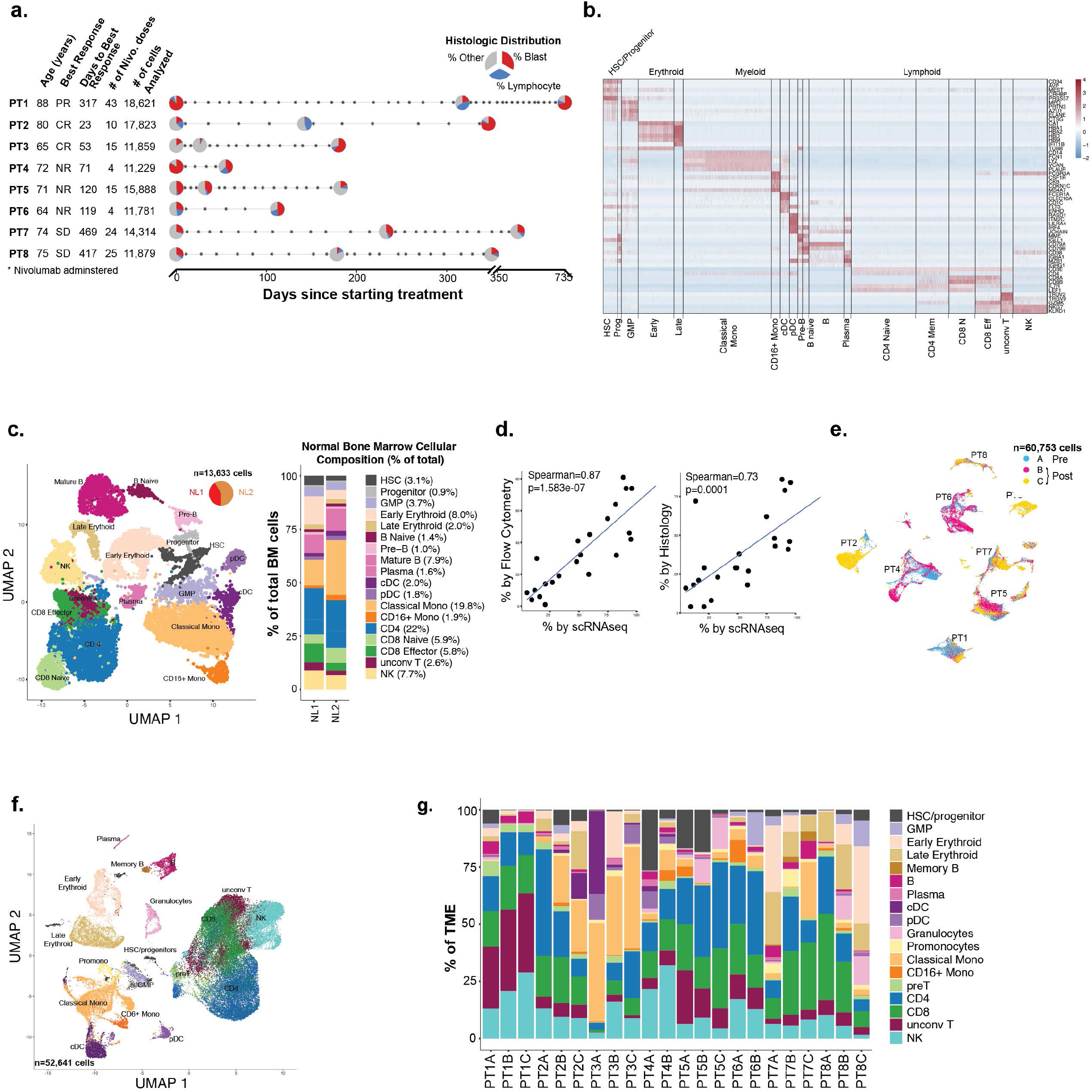
(a) Clinical design summarizing the age, response type per ELN, time to response, treatment frequency and the number of cells analyzed per patient. (b) Canonical gene expression markers to define the healthy bone marrow (BM) and tumor microenvironment (TME) cellular subsets. (c) UMAP-based trajectory analysis of healthy BM cellular components with frequency of each cell type. (d) Correlation between the number of cells detected by scRNA versus flow cytometry and histopathology. (e) UMAP clustering of AML cells. (f) UMAP clustering of TME components. (g) Distribution of TME components in AML patients at different timepoints (A is pre-treatment, B and C are post-treatment). Abbreviations, PR=partial response; CR=complete response; NR=no response; SD=stable disease. HSC=hematopoietic stem cell; GMP=granulocyte-monocytic progenitor; cDC=conventional dendritic cell; pDC=plasmacytoid dendritic cell; unconv T=unconventional T; NK=natural killer

### Cluster definitions in healthy and R/R AML BMs

To guide TME cluster definitions from AML cases, we first generated a BM cell atlas of 13,633 cells from 2 healthy BM donors and utilized canonical hematopoietic and immune gene expression markers to functionally annotate clusters(25–29) (Fig. 1b, Methods). Monocle3-based trajectory analysis(30–32) of healthy BM cells demonstrated a differentiation spectrum originating from hematopoietic stem cells (HSCs) and progenitors, leading to mature differentiated cells (Fig. 1c) consistent with previous reports(25,33). Also, AML cells demonstrated CD34 expression which was confirmed by validated multiparametric cytometry and/or immunohistochemistry (Supplementary Figure 2a-d), and clustered separately from TME components confirming a distinct transcriptional program (Supplementary Figure 3a-b), thus facilitating unambiguous downstream analyses.

A total of 60,753 AML and 52,641 TME cells from the 22 R/R AML BM aspirates from 8 patients passed quality check. The proportion of AML cells in the BM identified using scRNAseq closely correlated with proportion of AML blasts measured via flow cytometry (r=0.87, p=1.5×10^−7^) and histopathology (r=0.73, p=0.0001) on the clinical multiparametric flow-cytometry and histopathologic BM analysis done at the same clinical timepoint (Fig. 1d) providing strong validation for further analyses. Pre- and post-treatment AML cells clustered by patient (Fig. 1e). TME components from different patients clustered together and had different distributions (Fig. 1f-g). The clustering patterns of AML and TME cells were similar to other cancers and demonstrated significant intertumoral heterogeneity(23,25,34–36).

### Lineage tracing of R/R AML cells

To determine the likely origin of AML cells, we assessed their similarity to healthy BM cells by correlating the expression profile of each R/R AML cell with healthy BM cells and projected the result into a 2-dimensional trajectory map, similar to previously described(25). We measured the accuracy of the projection by randomly selecting 1000 cells from each predefined major TME component (HSC/GMP, T lymphocytes, B lymphocytes, monocytes, and erythroid cells) and projected it onto the healthy BM components. More than 85% of TME cells projected to the corresponding healthy BM cellular compartments which validated our approach (Supplementary Figure 4a). A median of 95.8% malignant cells from 21/22 (95%) timepoints projected onto HSCs of healthy BMs (Fig. 2a). After treatment, one patient (patient 4 (PT4), non-responder) demonstrated two dominant clusters: one cluster had high expression of CD34 and projected onto HSCs, and another cluster that did not express CD34 projected onto immunophenotypically committed monocytes (Fig 2b-c). These data demonstrated a HSC-like origin of R/R AML cells in all patients that persistent throughout treatment, while 1 patient (PT4, NR) also had a committed, monocytic phenotype emerging following treatment.

**Figure 2.**
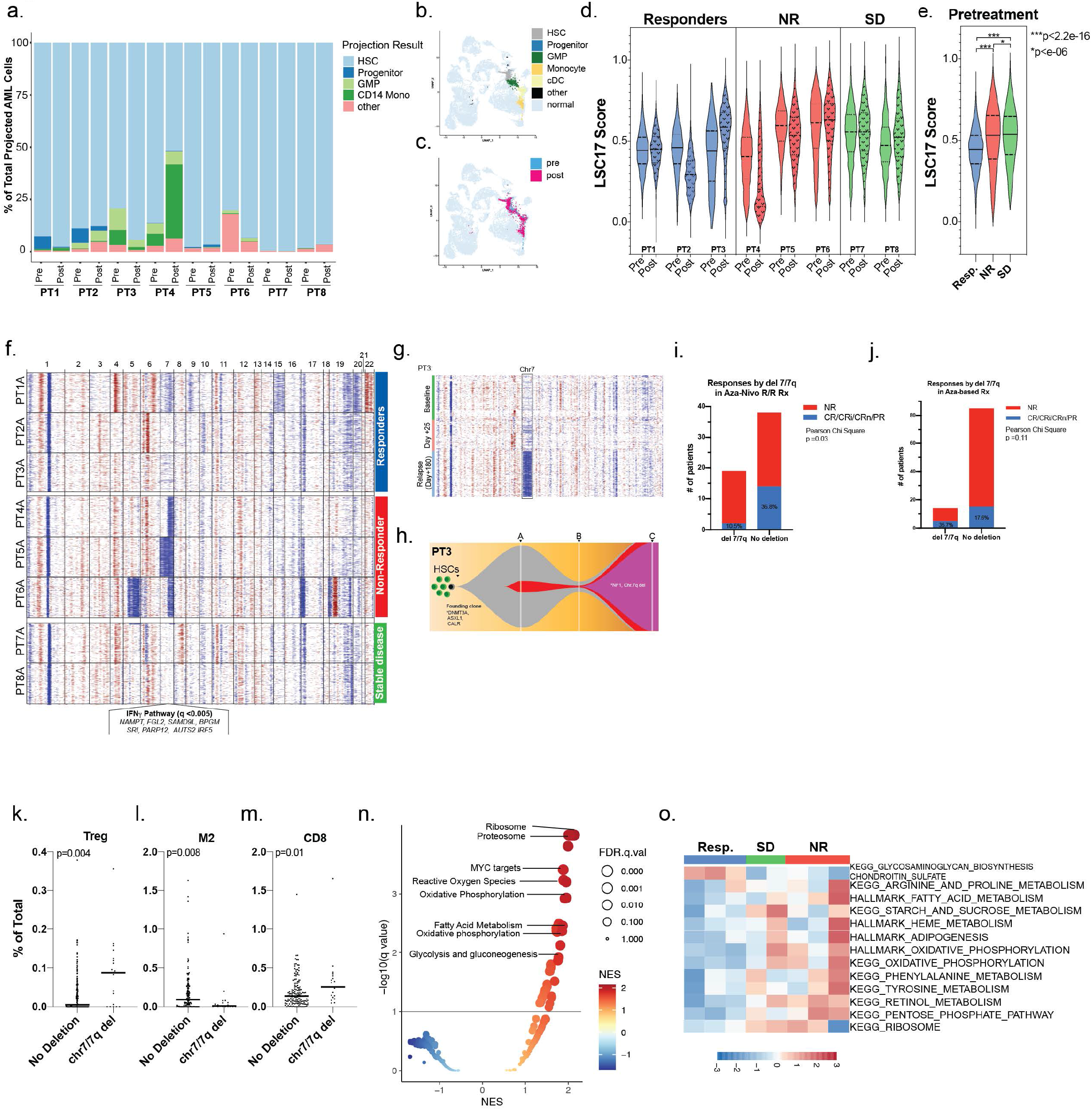
(a) Projection analysis of AML cells onto healthy BM cells. (b) PT4 (non-responder) projection UMAP onto healthy BM cells by projected cell type and (c) by treatment timepoint. (d) Violin plot of LSC17 scores in response groups and (e) at pretreatment. (f) Inferred copy number variation of representative 300 cells per patient at pretreatment (timepoint A). (g) Inferred copy number variation of the 3 timepoints (A, B and C) for PT3 (responder). (h) Fish plot of mutational evolution of PT3. (i) Correlation analysis for responders to azacitidine/nivolumab and (j) azacitidine-based therapy based on chromosome 7/7q deletion. (km): CIBERSORT analysis of immune cell components in AML from TCGA. (n) Gene set enrichment analysis of differentially expressed genes between responders and non-responders at pretreatment, (o) Heatmap of gene set variation analysis for metabolic and oxidative phosphorylation pathways at pretreatment.

Since resistance to chemotherapy and relapses in leukemia appear to be associated with leukemia stem cells (LSCs)(37), we utilized the LSC17 score that enriches for LSCs and correlates with treatment response and prognosis(38). Of the 17 genes in the LSC17 signature, 16 genes were detected in our scRNAseq profiling and were used for stemness scoring. The most significant change following treatment was noted in PT4 (NR) who demonstrated significantly decreased LSC17 stemness scores, which was consistent with detection of a committed monocytic lineage upon relapse (Fig. 2d). In an aggregate analysis of all cells at pretreatment timepoint, the 3 responders had lower LSC17 stemness scores compared to NR (p<2.2×10^−16^) and SD (p<10-6) patients (Fig. 2e). Thus, increased stemness appeared to be associated with resistance to ICB-based therapy.

### Chromosome 7/7q deletion is associated with resistance to ICB-based therapy

Since there are distinctive and recurrent copy number changes associated with AML, we utilized *inferCNV* tool to deduce large-scale copy number variations (CNVs) in scRNAseq expression data of AML cells (Methods). Our inferred CNVs were consistent with available clinical cytogenetics data for these patients from around the time of BM sampling (Supplementary Table 1). At pretreatment timepoint, 3/3 NR patients (PT4, PT5 and PT6) had inferred loss of chromosome 7/7q in most of their malignant cells (Fig. 2f). Of note, routine karyotyping of patient 1 (PT1; responder) demonstrated a complex profile with 7/19 cells harboring chromosome 7q deletion (Supplementary Table 1). This was congruent with the inferred intermediate chromosome 7q loss in our single cell RNA analysis (Fig. 2f). Interestingly, PT3 responder with CR) had an emergent chromosome 7q deletion (in 15/20 cells on karyotype), which preceded the clinical relapse (Fig. 2g-h). To further explore whether chromosome 7/7q was associated with resistance to ICB-based therapy, we evaluated 57 R/R AML patients treated on protocol NCT02397720 with azacitidine and nivolumab who had evaluable pretreatment cytogenetic profiling. Interestingly, only 10.5% (2/19) patients with chromosome 7/7q deletion achieved a CR/CRI/PR to treatment compared with 36.8% (14/38) of patients without deletion (p=0.03) (Fig 2i). To decouple azacitidine from nivolumab effect, we conducted similar analysis on an independent cohort of R/R AML cohort (n=99) treated on azacitidine-based clinical trials without ICB therapies at our institution. 35% (5/14) of patients with chromosome del 7/7q achieved a response (CR/CRI/CRn/PR) compared with 17.6% (15/85) without the deletion (p=0.11) (Fig. 2j). These data suggest that 7/7q loss is associated with resistance to the combination of nivolumab/azacitidine.

To investigate dysregulated molecular process and biological pathways in association with chromosome 7/7q resistance to ICB-based therapy, we conducted gene set enrichment analysis (GSEA) on chromosome 7q genes that were detected in our scRNA profiling. Interestingly, IFNγ pathway genes were significantly enriched (q<0.0005) in chromosome 7q region suggesting that IFNγ pathway loss may modulate resistance to ICB based therapies in AML, similar to previous findings in melanoma(20). We further estimated the abundance of the immune cells in AML cohort of TCGA in correlation with chromosome 7/7q loss via CIBERSORT(28) to investigate whether patients with chromosome 7/7q deletion had a deranged immune profile compared to patients without the deletion. Interestingly, T_reg_ (p= 0.004) cells were significantly higher in AML patients with chromosome 7/7q loss (n=19) compared to those with intact chromosome 7/7q (n=152) (Fig. 2k). Of note, M2 and CD8+ T cells proportion was also higher in patients with chromosome 7/7q deletion, however CIBERSORT does not allow further phenotypic description of CD8+ subsets (Fig. 2l-m). Additionally, TCGA AML patients with chromosome 7/7q loss had significantly worse survival than patients without chromosome 7/7q alteration (p=0.015) (Supplementary Figure 5a). Thus, chromosome 7/7q loss appears to harbor an immunosuppressive environment enriched in T_reg_ immune cells and confers inferior outcomes.

### Resistance to ICB-based therapy associated with higher oxidative phosphorylation and metabolic derangement in AML cells

We further explored potential tumor-intrinsic mechanisms by applying GSEA (Hallmark and KEGG gene sets) on differentially expressed genes between responders and NR in pretreatment AML cells. The highest enriched pathways in pretreatment NRs BM AML cells included oxidative phosphorylation, reactive oxygen species, and the metabolic pathways involving fatty acid metabolism and glycolysis/gluconeogenesis (Fig. 2n). Further, ribosomal pathway and *MYC* targets were significantly upregulated in non-responders BM AML cells (Fig. 2n), consistent with the role of MYC as a major regulator of ribosome biogenesis(39). To assess whether these pathways are similarly enriched in patients from the same response groups and not skewed by cell number contribution per patient, we applied gene set variation analysis (GSVA) to estimate the aforementioned pathway scores per patient(40). Responders had lower metabolic, oxidative stress, and ribosome biogenesis pathway activities at pretreatment compared to SD and NR patients (Fig. 2o). However, post-versus pre-treatment pathway GSEA for AML cells per patient did not reveal a discernible pattern that correlated with response groups (Supplementary Figure 6a). For instance, 6 or more patients had positive enrichment in at least one inflammatory or immune pathway activation at post-compared to pre-treatment (Supplementary Figure 6a). This suggested that while ICB-based therapy elicited activation of inflammatory pathways in AML cells, these were not sufficient to induce clinical responses. We therefore examined whether the T cell components of AML TME could shape responses to ICB-based therapy.

### T cell heterogeneity in R/R AML patients

We characterized the T cells in AML BMs as these are the main mediators of anti-tumor activity in response to ICB. We identified 5 distinct (2 classical and 3 non-classical) T cell phenotypes in 25,798 T cells from pre- and post-treatment timepoints in the BMs from these 8 patients (Fig. 3a, Methods). The 2 classical phenotypes were CD4+ and CD8+ cells, constituting 53% and 35% of the T-cells in the pretreatment BMs, and 30.9% and 37.4% of the T cells in the posttreatment BMs, respectively. The CD4:CD8 ratio of 1.51 in pretreatment BMs was lower than that in healthy donor BMs (1.88), and then decreased further to 0.82 following treatment. The 3 non-classical T cell phenotypes were gamma-delta (γδ) cells, mucosal associated invariant T-cells (MAIT) cells and unconventional T (unconv T) cells, constituting 2.3%, 2.1% and 7% of T cells in pretreatment BMs, versus 8.5%, 14.4% and 8.5% of T cells in posttreatment BMs, respectively. Thus, there was a significant increase in the proportion of CD8+, γδ and MAIT cells, while the proportion of CD4+ cells decreased following treatment on aggregate across all patients (Fig. 3a). Further, there were marked variations in the distribution of the five T cell phenotypes among the 8 patients and at different timepoints of treatment (Fig. 3b). At pretreatment, CD4+ cells were the most common cell type in responders (64.3%) and NR (42.48%) patients, whereas CD8+ cells were the most common in SD (53.48%) patients (Supplementary Figure 7a-c). Following treatment, CD8+ cells were the most common cell type in responders (30.28%) and SD patients (57.5%), whereas NR patients had persistently elevated CD4+ cells (52.91%) (Supplementary Figure 7d-f). Further, γδ and MAIT cells increased in responders following treatment, although this effect was primarily driven by from PT1 (responder, PR) (Supplementary Figure 7d). These findings demonstrated the TME heterogeneity within response groups, and dynamic changes occurring longitudinally for each patient.

**Figure 3.**
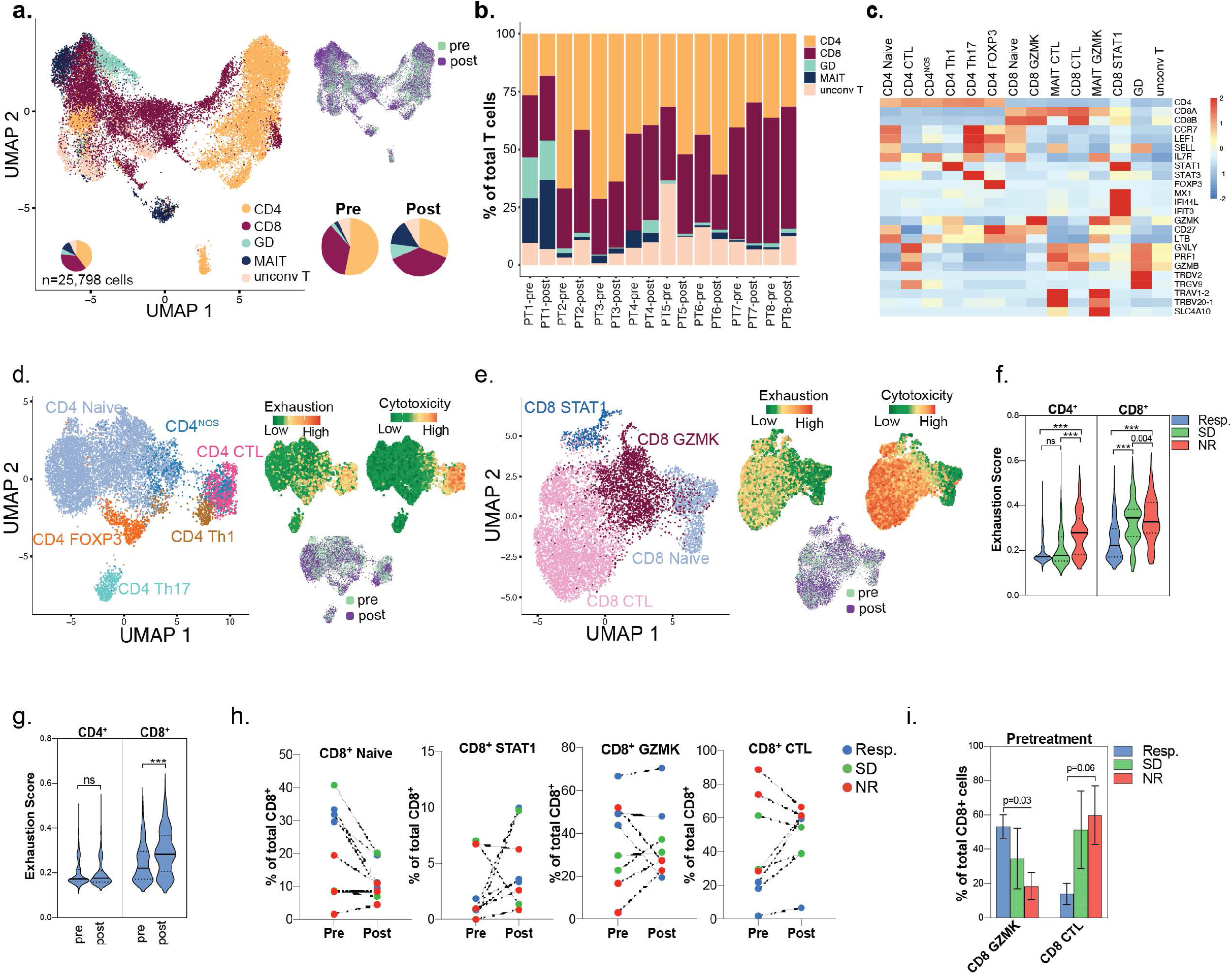
(a) UMAP of T cell subsets (b) Distribution of T cell subsets at different treatment timepoints. (c) Heatmap of canonical marker expression of identified T cell subsets. (d) UMAP of the different CD4 and (e) CD8 phenotypes with exhaustion and cytotoxicity scores projected onto the UMAP. (f) Exhaustion scores of CD4 and CD8 of different response groups at pretreatment. (g) Exhaustion score of CD4 and CD8 cells at pre and post treatment. (h) Pre- and post-treatment distribution change in CD8 subsets. (i) Pretreatment levels of CD8 GZMK and CD8 CTL in different response groups.

### CD8+ GZMK cells are higher, while CD8+ cytotoxic cells are lower in responders at pretreatment

We further classified the T cells based on canonical gene expression profile (Fig. 3c). We also measured the cytotoxic and exhaustion scores of these cells by utilizing GSVA for curated genes associated with these cell states(18,40,41). CD4+ cells clustered into CD4+ naïve and CD4+ effector subsets including FOXP3+ (T_reg_), T helper 1 cells (T_H_1), T_H_17 cells, and CD4+ cytotoxic (CTL) cells (Fig. 3d). Of note, a subset of CD4+ cells had no distinct gene expression profile and hereafter referred to as *not otherwise specified* (CD4_NOS_) (Fig. 3d, Methods). CD8+ clusters included CD8+ naïve, CD8+ STAT1 (enriched for STAT1 and expressed IFNγ pathway genes), CD8+ GZMK (express GZMK), and CD8+ CTL (expressing cytotoxic markers GZMB, GNLY and PRF1, but low expression of GZMK) (Fig. 3e, Methods). Exhaustion scores in CD4+ and CD8+ cells increased with increasing cytotoxic scores, while pre- and post-treatment cells clustered together by cell phenotype (Fig. 3d-e). At pretreatment timepoint, exhaustion scores of CD4+ and CD8+ cells were lowest in the 3 responders compared to the 2 SD and 3 NR patients (p<0.0001) (Fig. 3f). Following treatment, exhaustion scores of CD4+ were unchanged, while those of CD8+ significantly increased in the 3 responders (Fig. 3g).

There were no discernible patterns for the changes in the CD4+ subsets at pre- or posttreatment across response groups (Supplementary Figure 7g). However, the relative fraction of CD8+ naïve cells (among all CD8+ T cells) decreased following treatment in 7/8 patients, while 1 patient (non-responder) had a marginal increase (+2.9%) in CD8+ naïve cells (Fig. 3h). We noted that CD8+ GZMK constituted the majority of CD8+ cells in the 3 responders at pretreatment and were significantly more abundant in responders compared to NR (mean of 53.2% in responders 18.8% in NR, p=0.03) (Fig. 3i). Interestingly, at pretreatment timepoint, CD8+ CTL cells were the least abundant cells in the 3 responders and were lower compared to the 3 nonresponders (mean of 13.9% in responders vs 59.9% in NR, p=0.06) (Fig. 3i). Following treatment, all 3 responders had an increase in CD8+ CTL cells by an average of 2.8 folds (range: 2.1-3.6 fold), whereas 1 responder had 0.44 fold decrease in CD8+ GZMK cells, and another 2 responders had no notable change in CD8+ GZMK cells (0.9 and 1.05 fold). These data suggested more dynamic changes in the CD8+ than CD4+ subsets following ICB-based treatment in AML especially in responders, with the most remarkable differences occurring in the pretreatment CD8+ GZMK and CD8+ CTL components, which warranted further characterization to define the GZMK expressing cells in the context of AML.

### GZMK expressing CD8+ T cells intermediates naïve and cytotoxic cells

There are 5 human granzyme genes (*GZMA, GZMB, GZMH, GZMM*, and *GZMK*) with the function of only GZMA and GZMB well described(42). We observed high expression of GZMK in one of the CD8+ cell subsets, while CD8+ CTL cells were high in GZMB expression (Fig 3e and Supp Fig). To investigate whether the expression of granzymes could reflect a distinctive marker for cell state program, we applied Monocle3 pseudotemporal trajectory analysis and revealed that CD8+ GZMK are intermediary to CD8+ naïve and CD8+ CTL cells (Fig. 4a-b). We also observed distinctive expression of granzymes A, B, and K among the pseudotemporal axis of CD8+ cells. Specifically, GZMA was expressed ubiquitously in non-naïve CD8+ cells, whereas GZMB and GZMK expression profiles were largely distinctive to different cellular populations (Fig 4 c-e). Specifically, GZMB was expressed at the end of the pseudotemporal trajectory in the cytotoxic T cells, while GZMK expression was prominent in the intermediary cells. Further, the expression levels of the main cytotoxic gene *GNLY*, also delivers granzyme genes into target cells for effector functions(42), was significantly diminished in GZMK-expressing CD8+ cells compared to CD8+ CTL cells (fig. 4f). These findings were consistent with the lower cytotoxic and exhaustion scores in CD8+ GZMK cells compared to GNLY/GZMB-enriched CD8+ CTL cells (Fig. 3e, Supplementary Figure 7h). Interestingly, the co-stimulatory genes *LTB* and *CD27*, the stem-like T cell transcription factor *TCF7*, and the T cell memory transcription factor *EOMES* were highly expressed in CD8+ GZMK cells, but not CD8+ CTL cells (Fig. 4f). These findings, supported by pseudotemporal trajectory analysis, suggested a continuum of CD8+ cells with an intermediary, distinctive, CD8+ GZMK population in the TME of AML cells that is characterized by high GZMK expression, and harbored stem-like and memory T cell properties.

**Figure 4.**
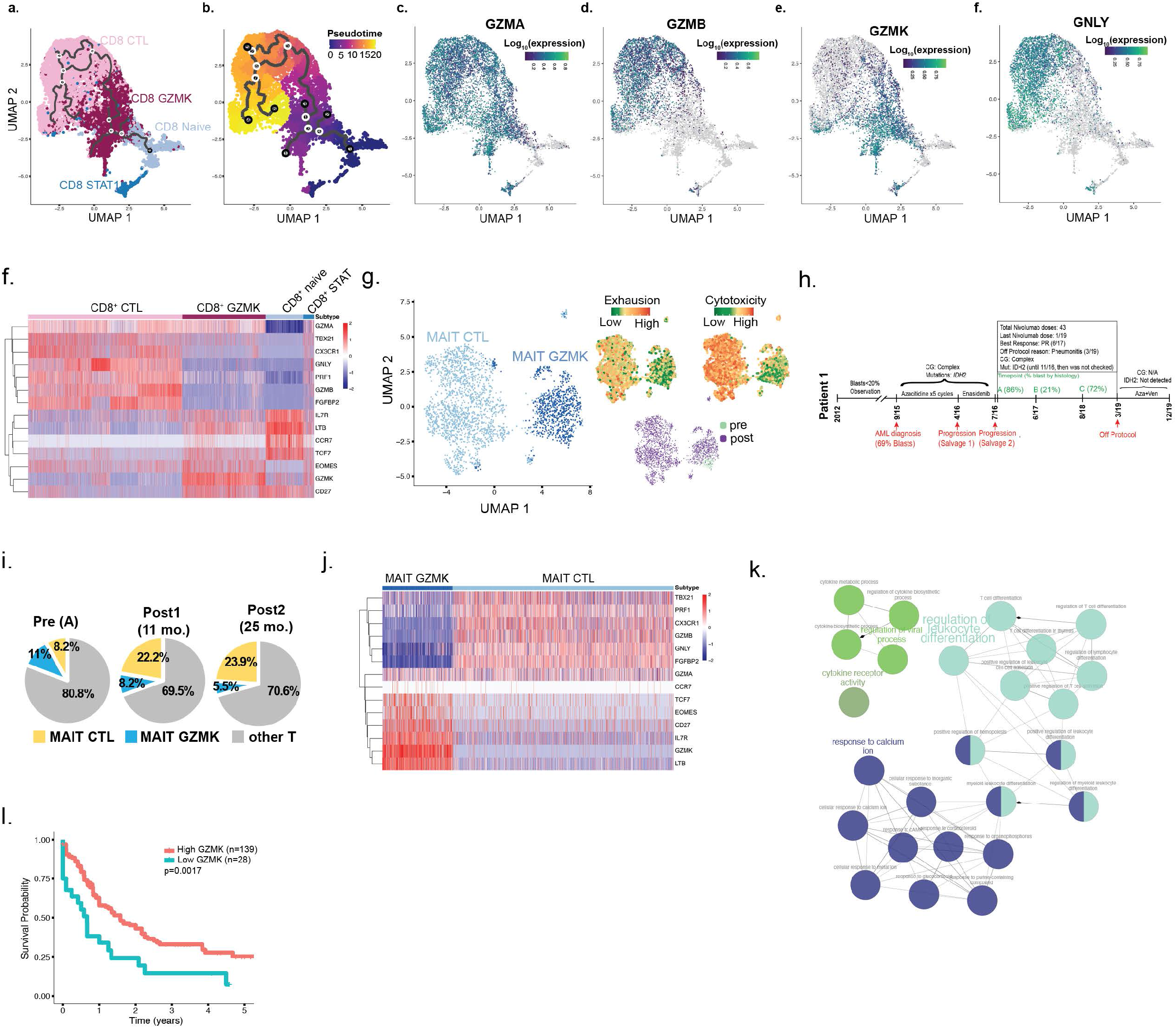
(a-b) Monocle3-based pseudotemporal analysis of CD8 subsets. (c-f) Expression of GZMA, GZMB, GZMK and GNLY in cells projected onto the trajectory of CD8 continuum. (f) Heatmap of differentially expressed genes between CD8 subsets. (g) UMAP of MAIT cells with exhaustion and cytotoxicity scores projection. (h) Clinical course of PT1 (responder). (i) Frequency of MAIT cells at pre and post treatment timepoints. (j) Heatmap of differentially expressed genes between MAIT subsets. (k) ClueGo plot of pathways enriched in GZMK-expressing cells. (l) Overall survival of TCGA AML patients based on GZMK expression.

### GZMK Expression Distinguishes MAIT Subsets in PT1 (responder, CR)

Interestingly, unbiased clustering of MAIT cells also revealed 2 distinct phenotypes: one enriched for less exhausted, GZMK expressing cluster (MAIT GZMK) and another enriched for GNLY/GZMB cytotoxic genes (MAIT CTL), similar to CD8+ cells (Fig. 4g). Of note, 89.9% of MAIT cells in our analysis were contributed by PT1 (responder) who had a unique clinical course (Fig. 4h). Briefly, PT1 had refractory AML to azacitidine (9/2015-4/2016), and to salvage with enasidenib (4/2016-07/2016) for *IDH2*-mutated refractory AML. At 88 years of age, he started a second salvage regimen with combined azacitidine/nivolumab with a partial response attained at 11 months from treatment initiation. He had sustained clinical benefit while on ICB-based therapy for 32 months until he developed ICB-induced pneumonitis necessitating switching treatment. Since this patient did not have a response to single agent azacitidine, then demonstrated a durable partial response to azacitidine/nivolumab, we postulate that the response was primarily driven by nivolumab. In PT1, the proportion of MAIT GZMK in TME decreased from 11%, to 8.2% then 5.5%, while MAIT CTL increased from 8.2%, to 22.2% to 23.9% following treatment (Fig. 2i), similar to the increase in CD8+ CTL cells we noted following treatment in responders.

Similar to CD8+ GZMK cells, MAIT GZMK cells were enriched for CD27, LTB, TCF7, and EOMES (Fig. 4j). Therefore, GZMK expression distinctly delineated subsets of CD8+ and MAIT cells suggesting a unique transcriptional program correlated with GZMK expression. We therefore conducted gene expression profiling comparing GZMK and CTL subsets of each of CD8+ and MAIT cells. Pathway enrichment of the 33 overlapping genes in the CD8+ and MAIT GZMK versus CTL signatures demonstrated highest enrichment for pathways involved in leukocyte differentiation, calcium signaling, and cytokine production (fig. 4k). Importantly, calcium signaling regulates T cell differentiation and activation, is required for achieving T cell functional specificity, and regulates cytokine secretion and cytotoxic pathways(43). These findings supported by pseudotemporal analysis and the pathway enrichment suggested an intermediary role for GZMK-enriched CD8 and MAIT cells in the lymphocyte spectrum. Interestingly, patients with higher GZMK (p=0.0017) expression had improved overall survival in AML TCGA data suggesting that inherent immunity with elevated GZMK can elicit improved outcomes (Fig. 4l).

### Higher T cell clonal expansion in responders and stable disease patients following ICB-based treatment

To determine the clonal relationship among different T cell subsets in response to treatment, we conducted single cell profiling of *α* and β TCR sequences from 4,742 and 26,095 T cells from healthy and R/R AML BMs, respectively. Only 7.2% (345/4742) of TCR clonotypes in healthy BMs were shared in 2 or more T cells, compared to 51% of TCR clonotypes in R/R AML. Further, the clonotype size in healthy BMs ranged from 1 to 16 TCR clonotypes, whereas that of R/R AML ranged from 1 to 1200 TCR clonotypes (Fig. 5a). This indicated that T cells in the TME of AML patients have less clonal diversity and more oligoclonal dominance compared to healthy BMs.

**Figure 5.**
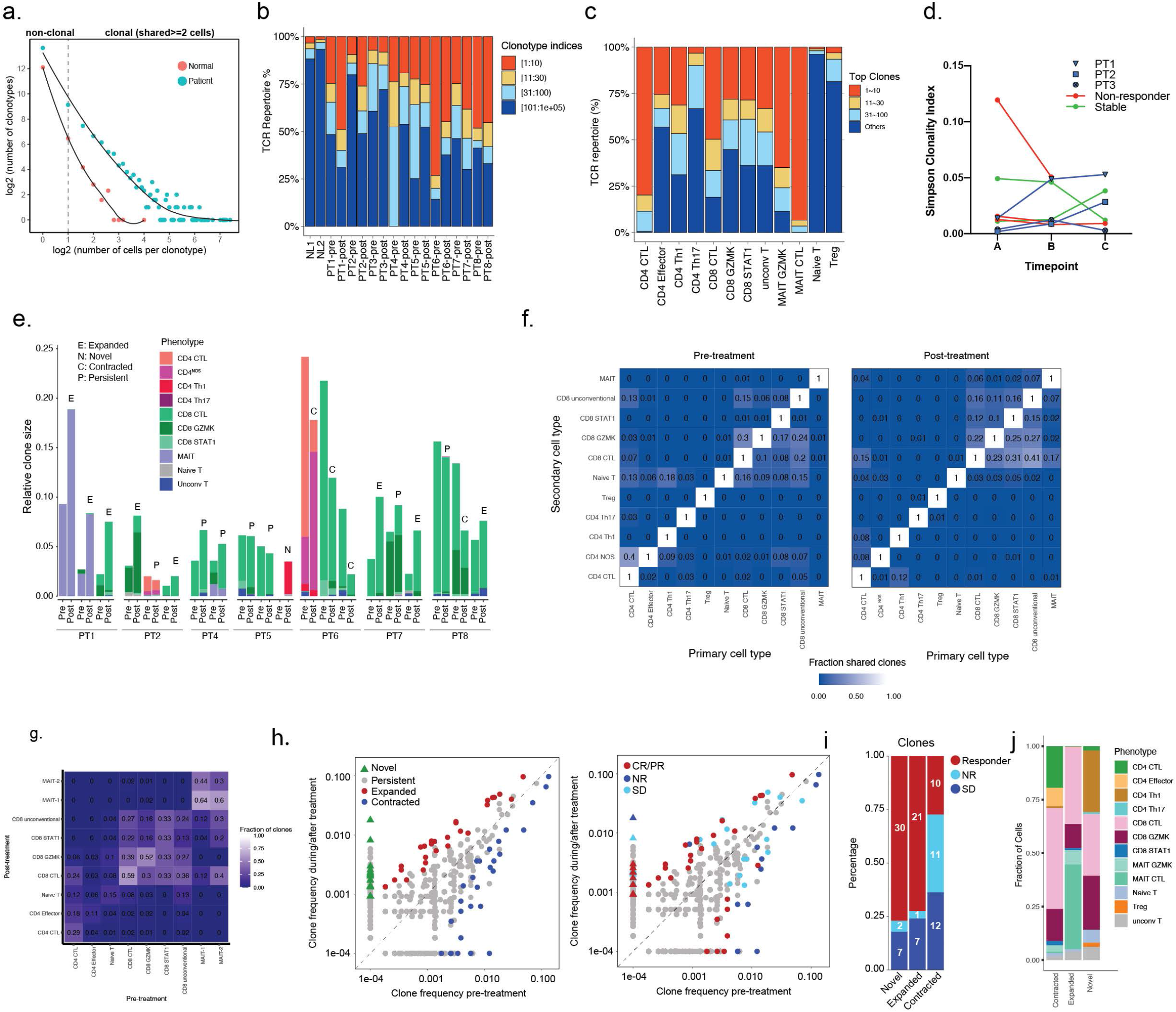
(a) Scatterplot of the correlation between the number of T cell clonotypes and the size of the clonotype. (b) Distribution of the most abundant clonotypes by patient and (c) cell type. (d) Simpson’s clonality index of individual patients at each of their respective timepoints (A is pretreatment and B and C are post-treatment). (e) Fraction of T cells from top clonotypes. (f) Heatmap of overlapping clonotypes between different celltypes at pre and post treatment timepoints. (g) Heatmap for observed phenotype transitions for matched clones. (h) Scatterplots of clonotypes change of pre-versus post-treatment and by response groups. (h) Number of novel, expanded and contracted clonotypes by response group and (j) cell type.

We next examined the degree of T cell clonal expansion that occurred longitudinally while on ICB-based therapy in patients with AML as well as in healthy BMs. As expected, the T cell repertoire from healthy BMs had few dominant clones (Fig. 5b). At pretreatment timepoint, there was marked variation in frequency of T cells contributing to the most abundant clonotype. Following treatment, 2/3 responders and 2/2 SD patients had expansion of their most abundant clonotypes (Fig. 5b). The third responder (PT3) had an increase in the most abundant clonotype following treatment initially, but then lost clonal dominance corresponding to the timing of acquisition of chromosome 7/7q loss, followed by disease progression. Conversely, all (3/3) NR patients had contraction of their most abundant clonotypes (Fig. 5b). Among T cells, cytotoxic subtypes had significant degree of clonal dominance (Fig. 5c). Based on Simpson clonality index, the 3 responders had an increase in their clonality following treatment (timepoint B) compared to pre-treatment (timepoint A) that persisted (timepoint C), except for PT3 (responder) who initially had an increase in clonality (timepoint B), then a subsequent decrease in clonality (timepoint C) corresponding with chromosome 7/7q that preceded his relapse (Fig. 5d). Three NR and 1 SD patients had relative reduction in their TCR clonality following treatment, while 1 SD patient (PT7) had a persistent increase in TCR clonality. This suggested that the expansion of dominant clonotypes is more likely to occur in responders to combined azacitidine/nivolumab treatment.

### Most abundant clonotype contributed by similar T cell phenotypes

We next evaluated the T cell phenotypes contributing to the top 3 most abundant clonotypes at pre- and post-treatment time points for each patient. The 3 most abundant clonotypes were largely contributed by cell types of same or similar phenotypes (Fig. 5e). For example, CD8+ GZMK and CD8+ CTL contributed to the most abundant clones of PT2, PT4, PT7 and PT8 (Fig. 5e). Also, MAIT GZMK and MAIT CTL of PT1 shared 2 of the most abundant clonotypes prior to treatment. Following treatment, the most abundant clonotypes expanded in 3/3 of the responders, but contracted in NR, while remaining unchanged in SD patients (Fig. 5e). Similar to what was seen in basal and squamous cell carcinoma following PD-1 treatment(23), there was no phenotypic instability of the dominants clonotypes in response to PD-1 blockade.

### Shared clonotypes among T cell lineages

We next explored whether the global clonotype profile within a T cell lineage (primary cell type) is shared with other T cell lineages (secondary cell type). The degree of shared clonotypes with the terminally effector CD8+ CTL cells increased following PD-1 blockade (Fig. 5f). To assess for clonotype stability, we evaluated the fraction of shared clones for each T cell phenotype between pre- and post-treatment timepoints (Fig. 5g). MAIT, CD8+ GZMK, and CD8+ CTL cells demonstrated relative clonotype stability, whereas CD4+ cells retained less clones following treatment (Fig. 5g). Further, 46% of pretreatment CD8+ GZMK clonotypes are shared with post-treatment CD8+ CTL. These findings suggest that there are shared clonotypes among closely related T cell lineages that expand following ICB therapy potentially.

### Novel and expanded clonotypes following treatment were predominantly from responders

We next evaluated the contraction, expansion, and emergence of novel clonotypes in response to treatment. Following treatment, novel clones represented 38.7% (39/101) of total clones, followed by contracted clones (32.6%) and expanded (28.7%) clones. However, 76.9% and 72.4% of novel and expanded clones were contributed by the responders. On the other hand, non-responders contributed only 5% and 3.4% of the novel and expanded clones, respectively (Fig. 5h-i). Similarly, among responders, the majority of clones were either novel or expanded, whereas NR had mostly contracted clones (Fig. 5i). CD8+ GZMK and CD8+ CTL contributed the most to novel clones, whereas MAIT CTL had the highest fraction of expanded clones, and CD8+ CTL had the highest fraction of contracted clones (Fig 5j). This indicated that the response to azacitidine/nivolumab is driven by emergence of novel clones and expansion of prior clones in CD8+ CTL and CD8+ GZMK cells, and in one patient, with a durable response, also from MAIT cells.

## Discussion

Disentangling AML cells from their complex, immune-rich microenvironment can reveal tumor intrinsic and extrinsic mechanisms that may have shaped resistance to ICB-based therapies and guide antileukemia therapeutic strategies. In this study, we leveraged single cell profiling of R/R AML BMs at pre- and post-treatment timepoints with azacitidine/nivolumab to determine tumor intrinsic and TME factors entwined with clinical responses. Our study revealed that AML cells harboring chromosome 7/7q loss, enriched for leukemia stem cell signature, and metabolic pathways involving oxidative stress were features associated with resistance to azacitidine/nivolumab therapy. Trajectory analysis revealed a continuous spectrum of CD8+ cells in the AML TME and unmasked a novel CD8+ subset with distinctively elevated GZMK expression harboring memory and stem-like characteristics, which was also a defining property in a subset of MAIT cells. Further, responses to treatment were associated with novel and expanded TCR clones. Since 7/8 patients in our study were refractory to prior hypomethylating agent treatment, we presume that the treatment effects observed are primarily driven via PD-1 inhibition. Our data suggest that in the appropriate context, T cells of the AML TME can be harnessed and activated to induce anti-leukemic activity.

While the number of patients who did not respond in this analysis is small, other studies have reported that deranged oxidative phosphorylation and heightened oxidative stress can impede T cell mediated cytotoxicity of tumor cells in melanoma(21,22,44,45). Similarly, oxidative phosphorylation signature is enriched in chemotherapy resistant AML and inhibition of BCL-2 targets oxidative phosphorylation to eradicate leukemia stem cells(46–48). The loss of chromosome 7/7q was correlated with loss of some of IFNγ genes in R/R AML and with immunosuppressive environment in *de novo* AML, which could confer resistance to immunotherapy as seen in other cancers(20,49,50). These findings are of particular importance as cytogenetic profiling, which is part of routine pathologic assessment for AML patients prior to treatment initiation, may potentially guide patient selection for ICB-based therapy trials in AML.

GZMB is well described in mediating apoptosis in target cells(51–54). However, GZMK is considered an ‘orphan’ granzyme due to its ambiguous function, with some studies considering it a cytolytic marker, while others suggesting no cytotoxic activity(35,42). We identified a continuum of CD8+ phenotypes that are distinguished by GZMB and GZMK expression, with increased exhaustion with increasing GZMB levels, and memory and stem-like features with increasing GZMK expression. Noteworthy, exhausted and cytotoxic lymphocytes can be difficult to distinguish without functional assays as computationally-derived markers of each can overlap. We hypothesize that the CD8+ naïve population maintains CD8+ GZMK pool via a stable influx, which ultimately differentiates into CD8+ CTL. This is supported by our pseudotemporal trajectory analysis of CD8+ phenotypes, ebbing of CD8+ naïve cells and the surge in exhausted CD8+ CTL cells following treatment, and the shared TCR repertoire between CD8+ GZMK and CD8+ CTL that emerged after PD-1 inhibition.

CD8+ GZMK cells in AML TME did not conform to previously reported granzyme-expressing CD8+ phenotypes. CD8+ GZMK had low expression of cytotoxic genes but harbored high expression of both transcription factors TCF7 and EOMES. TCF7-expressing CD8+ cells mediate and sustain immune responses via stem-like activities in a melanoma murine model treated with a PD-1 inhibitor, and following infection with lymphocytic choriomeningitis virus(55–57). Additionally, TCF7+ CD8+ T cells define stem-like T cells in cancer patients which have enhanced self-renewal, multipotency, and give rise to more terminally differentiated effector CD8+ T cells in tumors, further supporting our continuum model(58,59). EOMES enables enrichment of memory CD8+ T cells in the BM niche, promotes memory cell formation, and its expression in CD8+ cells correlates with response to immunotherapy in melanoma(22,60,61). Collectively, these findings reveal that CD8+ GZMK cells constitute a novel subset of memory CD8+ cells in the AML TME, enriched for stem-like markers, and are potentially the clonal origin for CD8+ CTL following PD-1 inhibition. Interestingly, the GZMK-expressing phenotype in T cells was also recapitulated in MAIT cells suggesting that GZMK is a *de facto* marker of cellular states in T cells. The exceptional long duration of response in 1 R/R AML patient (PT1) was associated with MAIT cell expansion, consistent with the role of MAIT cells in eliciting potent antitumor activities (62–64). To our knowledge, this is the first study that reports PD-1 blockade can induce MAIT cell expansion and response.

Importantly, the majority of the responses attained were primarily driven by either expanded or novel TCR clonotypes. Of these novel and expanded clonotypes, 75% were from responders, and approximately 20% from SD patients, while only 5% were detected from NR. Thus, the degree of novel or expanded clones corresponded with the degree of response. These findings demonstrate that PD-1 based therapy is capable of selectively expanding presumptive anti-tumor clones in the AML TME.

In summary, we report the largest number of AML BM cells profiled at single cell resolution in a single study allowing for deeper understanding of the AML TME in the context of ICB-based therapy. Our results provide insight into the intrinsic AML characteristics that could guide in negative selection of checkpoint-inhibition based trials in patients with chromosome 7/7q loss. Our results further validate approaches to combine inhibition of oxidative phosphorylation by venetoclax-based therapies with PD-1 inhibition in AML, currently being pursued in early phase clinical trials (NCT02397720, NCT04284787, NCT04277442). Our results demonstrate that the subverted T cells can be reinvigorated via PD-1 blockade and elicit responses in AML and warrants further functional characterization of GZMK expressing lymphocytes in mediating antileukemic responses. These data build a framework for deeper understanding of the molecular dynamics of ICB in leukemia that can hopefully drive strategies to develop optimal combinatorial approaches to improve efficacy and outcomes for these patients.

## Methods

### Human Subjects and treatment regimen

Patients >= 18 years of age who had failed prior therapy for AML were eligible to participate on combined azacitidine and nivolumab trial (ClinalTrials.gov identifier: NCT02397720; full protocol is included in Supplementary files). Bone marrows from 8 patients on NCT02397720 protocol and 2 healthy donors were characterized in this study. All patients had histologically proved relapsed or refractory acute myeloid leukemia. Treatment consisted of azacitidine 75 mg/m2 days 1 to 7 administered intravenously (i.v.) over 60 to 90 minutes or subcutaneously, and nivolumab 3mg/kg administered as a 60 to 90 minute i.v. infusion on days 1 and 14 of each cycle. Each cycle consisted of 28 days and were repeated every 4 to 6 weeks, depending on count recovery. Patients continued on therapy as long as they had evidence of clinical benefit. Response assessment was conducted using European Leukemia Network (ELN) response criteria. A written informed consent that was approved by the internal review board of University of Texas M D Anderson Cancer Center was obtained. The study was conducted in accordance with the Declaration of Helsinki.

### Sample collection and processing

Bone marrow (BM) biopsies were routinely collected prior to treatment initiation and at different timepoints for response assessments. BM samples were freshly frozen with 20% fetal calf serum (FCS) and 10% DMSI in DMEM media, and stored in liquid nitrogen.

### Cells preparation for single cell profiling

All BM samples stored in liquid nitrogen were retrieved right before sample processing. To maximize the cellular viability recovery, samples were processed in batches according to in house developed protocol and 10x Genomics “Demonstrated Protocol Cell Preparation Guide” (Document CG00053). Briefly, cells were gently thawed in water bath at 37°C until partially thawed and immediately placed on ice. Next, cells were gently transferred to a 10ml media (10ml alphaMEM + 20%FCS) and centrifuged (1500 rpm for 5 min). After removal of supernatant, the cell pellet was carefully resuspended in 10ml enriched media (alphaMEM+20% FCS supplemented with 500 μL Heparine, 15μL DNase and 500μL MgSO4), followed by incubation in 37°C for 15min. After incubation, cells were centrifuged and gently washed twice in 1.5-3mL of 0.04% BSA in PBS. Additionally, cells were passed through strainer (0.35um - 0.4um Flowmi Cell Strainer) to eliminate cell clumps. Next, cells were stained with 0.4% Trypan blue and quantified and assessed for viability using the cell automated counting machine Cellometer Mini (Nexcelom, Lawrence, MA, US), as well as using standard hemocytometer and light microscopy.

### Library preparation for 10x Genomics single-cell 5’ and VDJ sequencing

The scRNA-Seq and scTCR enriched libraries were prepared using the 10x Single Cell Immune Profiling Solution (https://www.10xgenomics.com/products/single-cell-immune-profiling/), according to the manufacturer’s protocol “CG000086 Chromium Single Cell V(D)J Reagent Kits User Guide Rev G (v1 Chemistry), (10x Genomics, Pleasanton, CA, USA)”. The brief description of the key steps conducted for libraries generation are summarized below.

#### cDNA generation from single cell

the single cell suspensions with a targeted cell recovery of 10,000 cells per sample were mixed with Master Mix and loaded into the Chromium Chip A along with the barcoded Single Cell VDJ 5’ Gel Beads v1 and Partitioning Oil. The nanoliter - scale Gel Beads-in-emulsion (GEMs) were generated using 10x Chromium Controller. Next, GEMs were captured and incubated (Step 1: 53°C for 45 min, Step 2: 85°C for 5 min) to produce 10x Barcoded with unique molecular index (UMI), full-length cDNA from polyadenylated mRNA in reverse transcription reaction. Following steps included breaking the GEMs, Post GEM-RT Cleanup and PCR amplification of released cDNA with primers against common 5’ and 3’ ends added during GEM-RT (Step 1: 98°C for 45 sec, Step 2: 98°C for 20 sec, Step 3: 67°C for 30 sec, Step 4: 72°C for 1 min, Step 5: 72°C for 1min, Hold: 4°C; Steps 2-4 were performed in total of 13 cycles). The obtained cDNA was cleaned-up using SPRIselect Reagent Kit (Beckman Coulter) and quantified using an Agilent 4200 Tape Station HS D5000 Assay (Agilent Technologies). Such prepared cDNA was further used to generate 5’ gene expression (5’GEX) libraries and TCR-enriched libraries.

#### Construction of 5’GEX libraries

50ng mass of cDNA of each sample was used. The main steps of 5’GEX library preparation included: (1) fragmentation, end repair and A-tailing followed by double sided size selection (2) adaptor ligation with post-ligation cleanup, and 3) sample index PCR followed by the double sided cleanup and QC. The enzymatic fragmentation and size selection (SPRIselect Reagent Kit, Beckman Coulter) were used to optimize the cDNA amplicon size followed by adding P5 and P7 sample indexes (used in Illumina sequencers), and Illumina R2 sequence via processes of end repair, A-tailing (Hold: 4°C, Step 1: 32°C for 5 min, Step 2: 65°C for 30 min, Hold: 4°C), adaptor ligation (20°C for 15 min, Hold: 4°C) and Sample Index PCR (Step 1: 98°C for 45 sec, Step 2: 98°C for 20 sec, Step 3: 54°C for 30 sec, Step 4: 72°C for 20 sec, Step 5: 72°C for 1min, Hold: 4°C; Steps 2-4 were performed in total of 14 cycles) using Chromium i7 Sample Index (PN-220103, 10x Genomics).

#### TCR enrichment step

5ul of cDNA of each sample was used. The TCR transcripts were enriched by two rounds of PCR amplification (Step 1: 98°C for 45 sec, Step 2: 98°C for 20 sec, Step 3: 67°C for 30 sec, Step 4: 72°C for 1 min, Step 5: 72°C for 1min, Hold: 4°C; Steps 2-4 were performed in total of 10 cycles) with primers specific to the TCR, with sample cleanup between each run of PCR. Noteworthy, the P5 was also added during enrichment. The double sided size selection was performed on enriched target followed by QC step.

#### Construction of TCR enriched libraries

50ng mass of TCR enriched samples were used. The main steps of the process included: (1) fragmentation, end repair and A-tailing (Hold: 4°C, Step 1: 32°C for 2 min, Step 2: 65°C for 30 min, Hold: 4°C), (2) adaptor ligation (20°C for 15 min, Hold: 4°C) with post-ligation cleanup, (3) sample index PCR amplification (Step 1: 98°C for 45 sec, Step 2: 98°C for 20 sec, Step 3: 54°C for 30 sec, Step 4: 72°C for 20 sec, Step 5: 72°C for 1min, Hold: 4°C; Steps 2-4 were performed in total of 9 cycles) using Chromium i7 Sample Index (PN-220103, 10x Genomics), followed by the double sided cleanup and QC step.

### Sequencing

All libraries were checked for the fragment size distribution using Agilent 4200 Tape Station HS D5000 Assay (Agilent Technologies) and quantified with Qubit Fluorometric dsDNA Quantification kit (Thermo Fisher Scientific, Waltham, MA, US). Each of 5’GEX and TCR enriched libraries were prepared with unique indexes allowing for multiplexing. The 5’GEX libraries were sequenced each of on a separate lane of HiSeq4000 flow cell (Illumina) to target sequencing depth of 50,000 read pairs per sample, in total of four batches. All TCR enriched libraries were equimolarly pooled and sequenced in one sequencing run using NovaSeq6000 S2-Xp 100 (Illumina) as required lower sequencing depth of 5,000 read pairs per sample. All libraries were sequenced at the ATGC MDACC core facility under 10x Genomics recommended cycling parameters for 5’GEX libraries (Read 1 - 26 cycles, Read 2 - 98 cycles; with few exceptions for samples sequenced on the same flow cell along with scATAC libraries: Read 1 - 100 cycles, Read 2 - 100 cycles) and for scTCR enriched libraries (Read 1 - 26 cycles, Read 2 - 91 cycles), with sequencing format of 100nt. All datasets generated during and/or analyzed during the current study will be available from the corresponding authors.

### Raw sequencing data processing, quality check, data filtering, doublets removal, batch effect evaluation and data normalization

**T**he raw scRNA-seq data were pre-processed (demultiplex cellular barcodes, read alignment, and generation of gene read count matrix) using Cell Ranger Single Cell Software Suite provided by 10x Genomics. Detailed QC metrics were generated and evaluated. Genes detected in fewer than 3 cells and cells with low complexity libraries (less than 200 genes were detected) were filtered out and excluded from subsequent analysis. Low-quality cells where >15% of transcripts derived from the mitochondria were considered apoptotic and also excluded. Following the initial clustering, we removed likely cell doublets from all clusters. Doublets were identified by following methods as previous described (65) 1) library complexity-cells are outliers in terms of library complexity. 2) Cluster distributiondoublets or multiplets likely form distinct clusters with hybrid expression features and exhibit an aberrantly high gene count; 3) cluster marker gene expression--cells of a cluster express markers from distinct lineages (e.g., cells in the T-cell cluster showed expression of epithelial cell markers; 4) Non-T cells expressing TCR or Non-B cells expressing BCR.

### Unsupervised cell clustering, dimensionality reduction using t-SNE and UMAP, and cluster relationship analysis

Library size normalization was performed in Seurat (66) on the filtered gene-cell matrix to obtain the normalized UMI count as previously described (67). Then, the normalized gene-cell matrix was used to identify highly variable genes for unsupervised cell clustering in Seurat. The elbow plot was generated with the ElbowPlot function of Seurat and based on which, the number of significant principal components (PCs) were determined. Different resolution parameters for unsupervised clustering were then examined in order to determine the optimal number of clusters. For visualization, the dimensionality was further reduced using Uniform Manifold Approximation and Projection (UMAP) (68) methods with *Seurat* function *RunUMAP*. In addition, *Monocle 3* alpha (http://cole-trapnell-lab.github.io/monocle-release/monocle3/)(69) was applied as an independent tool for unsupervised clustering analysis (function *cluster_cells*) and UMAP was used by default with the *Monocle* functions *reduce_dimension* and *plot_cells* for dimensionality reduction and visualization of the *Monocle* clustering results. *Monocle 3* alpha was also used to construct the single-cell trajectories. The function *learn_graph* was run with default parameters. Batch effects was corrected using the R package HARMONY before clustering analysis in Seurat and using the *Alignment* function in Monocle3 before trajectory analysis.

### Determination of major cell types and cell states

To define the major cell type of each single cell, differentially expressed genes (DEGs) were identified for each cell cluster using the *FindAllMarkers* analysis in the Seurat (66) package and the top 50 most significant DEGs were carefully reviewed. In parallel, feature plots were generated for top DEGs and a suggested set of canonical immune and stromal cell markers, a similar approach as previously described (70,71), followed by a manual review process. Enrichment of these markers (e.g. *PTPRC* for immune cells; *CD3D/E* for T cells; *CD8A/B* for CD8 T cells, *IL7R/CD4/CD40LG* for CD4 T cells; *CD19/MS4A1/CD79A* for B cells, *etc.*) in certain clusters was considered a strong indication of the clusters representing the corresponding cell types. The two approaches are combined to infer major cell types for each cell cluster according to the enrichment of marker genes and topranked DEGs in each cell cluster, as previously described (71).

### Infer large-scale copy number variations (CNVs)

The tool *inferCNV* (https://github.com/broadinstitute/inferCNV) was applied to infer the large-scale CNVs from scRNA-seq data and monocytes from normal bone marrow of this dataset were used as a control for CNV analysis. Initial CNVs were estimated by sorting the analyzed genes by their chromosomal locations and applying a moving average to the relative expression values, with a sliding window of 100 genes within each chromosome, as previously described (34,72). Finally, malignant cells were distinguished from normal cells based on information integrated from multiple sources including marker genes expression, inferred large-scale CNVs, and their cluster distribution per patient with cells from normal bone marrow.

### Pathway enrichment analysis

For pathway analysis, the curated gene sets (including Hallmark and KEGG gene sets) were downloaded from the Molecular Signature Database (MSigDB, http://software.broadinstitute.org/gsea/msigdb/index.jsp), single-sample GSVA (ssGSVA) was applied to the scRNA-seq data and pathway scores were calculated for each cell using *gsva* function in GSVA software package (40). Heatmaps were generated using the *heatmap* function in pheatmap R package for filtered DEGs.

### TCR V(D)J sequence assembly, clonotype calling, TCR diversity and clonality analysis and integration with scRNA-seq data

Cell Ranger v3.0.2 for V(D)J sequence assembly was applied for TCR/BCR reconstruction and paired clonotype calling. The CDR3 motif was located and the productivity was determined for each single cell. The clonotype landscape was then assessed and the clonal fraction of each identified clonotype was calculated. The TCR clonotype diversity matrix was calculated using the tcR R package (73). The clonotype data was then integrated with the T-cell phenotype data inferred from single cell gene expression analysis based on the shared cell barcodes.

### Immune cell deconvolution of TCGA sample

CIBERSORT (28) was applied to the normalized bulk RNA-seq data with the LM22 gene signature to estimate the relative fractions of 22 immune cell types.

### Statistical Analysis

Statistical differences were calculated with an unpaired Student’s t-test for 2-tails. Spearman’s correlation coefficient was used. A value of p<0.05 was considered statistically significant. For multiple t-tests, false discovery rate with q<0.05 was used for statistical significance. All statistical analysis were performed using packages in R (R Foundation for Statistical Computing) and in GraphPad Prism Software version 8.

## Acknowledgements

We would like to acknowledge the Advanced Technology Genomics Core (ATGC) at M D Anderson and 10xgenomics for sequencing and technical support. The work was funded in part by T32 NIH fellowship to H.A.A; CPRIT MIRA (RP160693) and Department of Leukemia SPORE (5 P50 CA100632-09) to S.M.K.; CA016672 and NIH 1 S10OD024977-01 to Advanced Technology Genomics Core (ATGC) and MDACC start-up to K.R.; MDACC startup to L.W.; CPRIT MIRA (RP160693) and MD Anderson Cancer Center Leukemia SPORE CA100632 to S.K; the MD Anderson Cancer Center Support Grant (CCSG) CA106672, the M D Anderson Cancer Leukemia SPORE CA100632, the Dick Clark Immunotherapy Fund and generous philanthropic contributions to the MD Anderson Moon Shots Program to ND; APOLLO, Welch Foundation and CPRIT grants to A.F..

## Author Contributions

Conceived the project: H.A.A., N.D. and A.F. Lead library preparation and experimental optimization: K.T., and K.R.. Lead computational analysis: D.H. and L.W. Optimized experiments, analyzed the data, wrote the manuscript: H.A.A. D.H., K.T., L.W., K.R., N.D. and A.F.. Contributed to data analysis, study design, experimental design, optimization and/or writing the manuscript: J.I.S., P.K.R., Z.A., W.W, R.W., F.W., G.A., M.R.G., K.T., L.T., J.Z., S.B.. Bone marrow processing and clinical annotations: G.A., J.T.M., Z.A., S.M.K.

Statistical tests and design: J.N. and M.D. Contributed conceptually to data analysis, writing and design: M.K., G.G.M., P.S. and J.P.A., S.M.K. Hematopathology: W.W.

---

**Supplementary Table 1.** Clinical characteristics of patients 1 to 8 on combined nivolumab/azacitidine study.

| Patient ID | Age | Prior HMA | Time to Response Assessment (Months) | Best Response | Off protocol reason | Pretreatment Cytogenetics |
| --- | --- | --- | --- | --- | --- | --- |
| PT1 | 88 | Yes | 10.57 | PR | Pneumonitis | 75~90<4n>,XXYY,-3,-7,-8,-9,+12,-13,-14,-16,+19,-20,+22,+22,+mar[cp13]/46,XY[7] |
| PT2 | 80 | Yes | 0.77 | CR | Relapse | 46,XY[20] |
| PT3 | 65 | No | 1.77 | CR | Relapse | 46,XY,del(12)(p12)[5]/46,XY[15] |
| PT4 | 72 | Yes | 2.37 | NR | No response | 46,XY,del(7)(q22)[20] |
| PT5 | 71 | Yes | 4.00 | NR | No response | 45,XY,-7[19]/46,XY[1] |
| PT6 | 64 | Yes | 3.97 | NR | No response | 45,XX,5q-,6p-,<br>-17,der(20)add(20q),+mar[3]/46,XX,-5,6p-,<br>-7,-17,19q+,der(20)add(20q),+2mar[7] |
| PT7 | 74 | Yes | 15.63 | SD | Loss of clinical benefit | 47,XY,+8[5]/47,idem,del(8)(q21.2q24.3)[8]/46,XY[7] |
| PT8 | 76 | Yes | 13.90 | SD | Loss of clinical benefit | 46,XY,t(3;14)(q21;q32)[1]/46,XY,-14,add(20)(q13.3),+mar[1]/46,XY[18] |

\*PR=partial response; CR=complete response; NR=no response; SD =stable disease

**Supplementary Figure 1.**
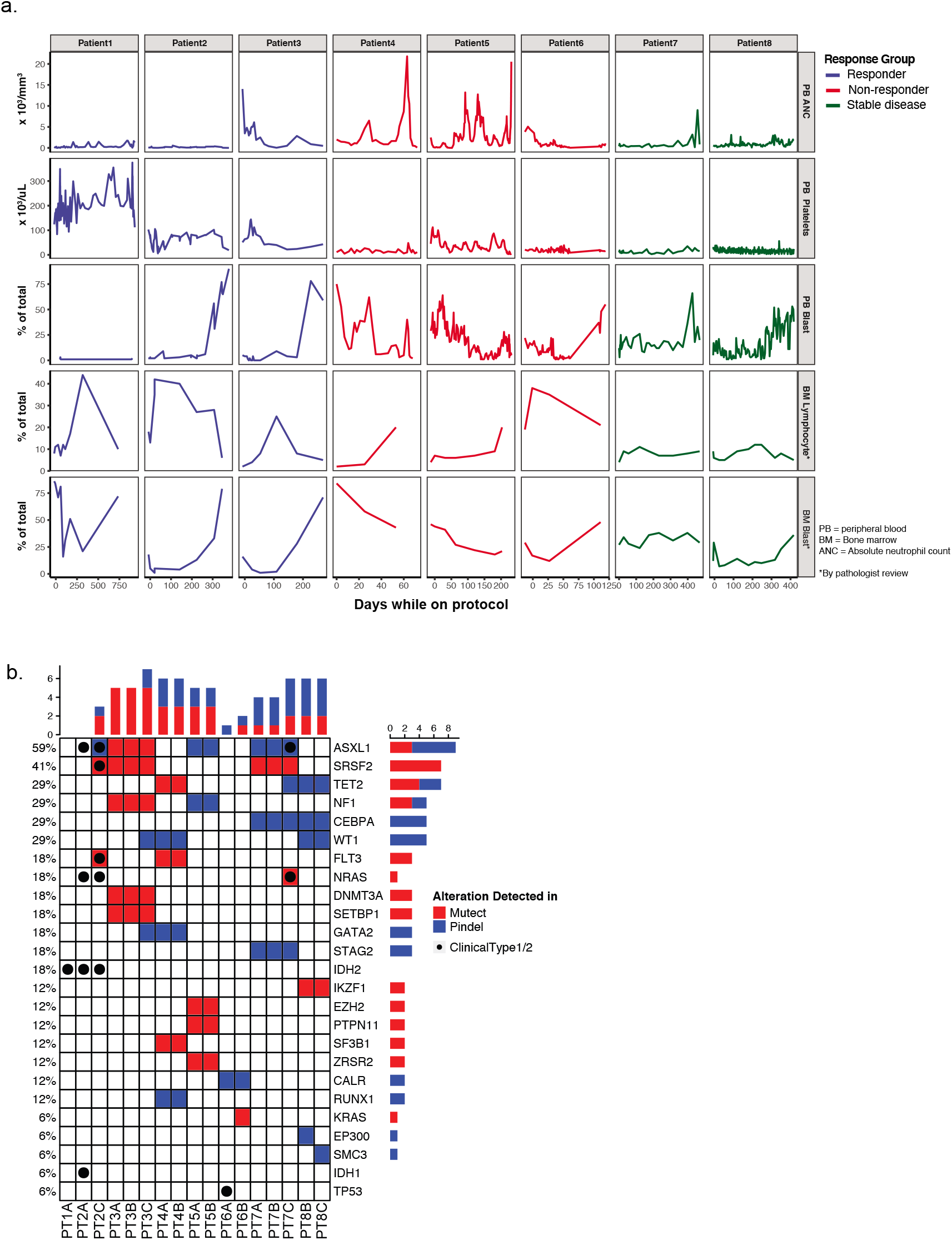
(a) Bone marrow (BM) and peripheral blood (PB) laboratory results while on azacitidine/nivolumab protocol. (b) Targeted DNA sequencing with cancer gene list (Takahashi et al Blood 2018) as well as a CLIA-certified molecular diagnostic assay at the different treatment timepoints when available.

**Supplementary Fig. 2.**
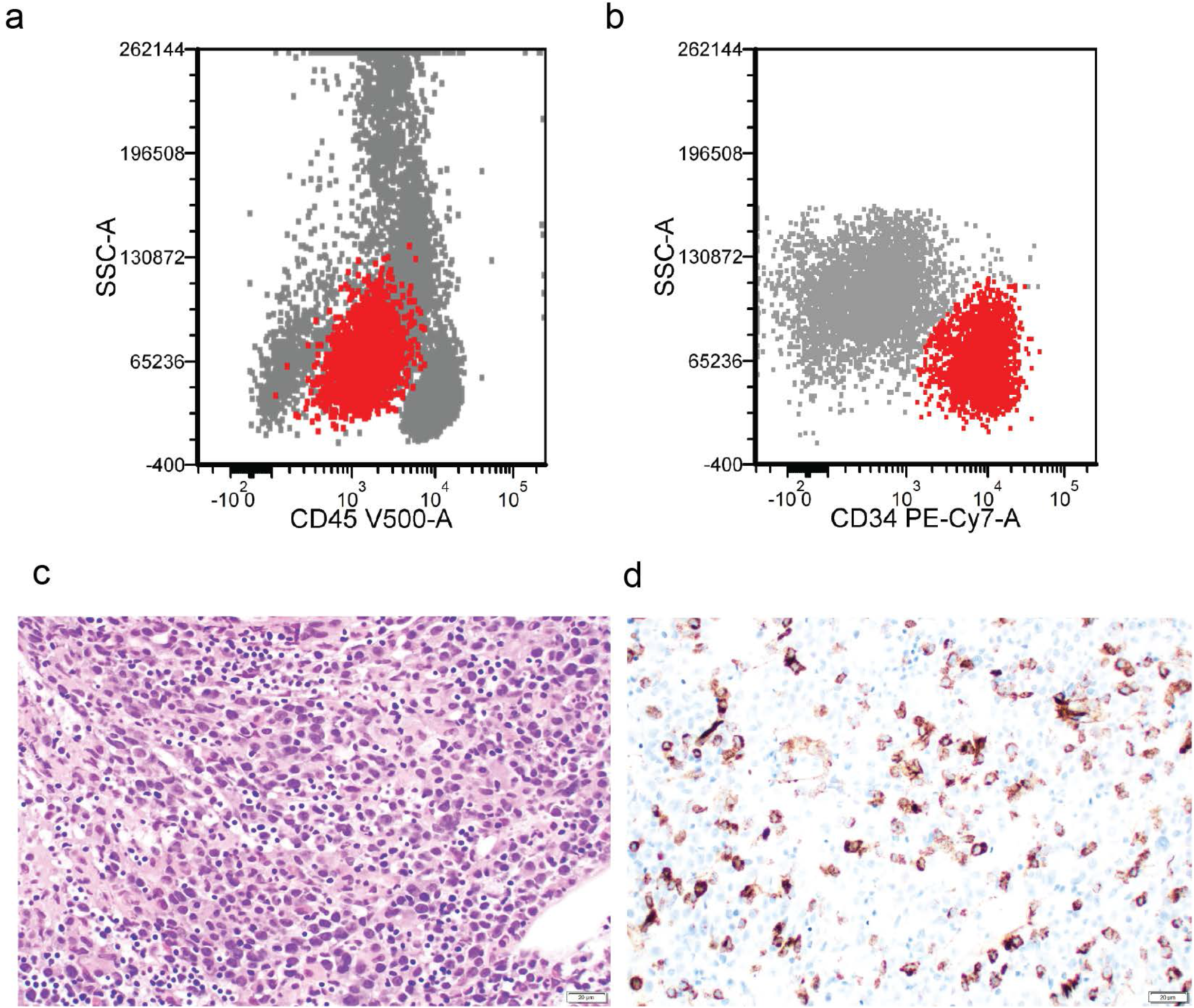
(a-b) Representative flow cytometry analysis from BM cells demonstrating CD34+ve blasts. (c) Hematoxylin and eosin staining of a representative BM biopsy. (d) Immunohistochemistry with CD34 antibody demonstrating CD34 positive blasts.

**Supplementary Fig. 3.**
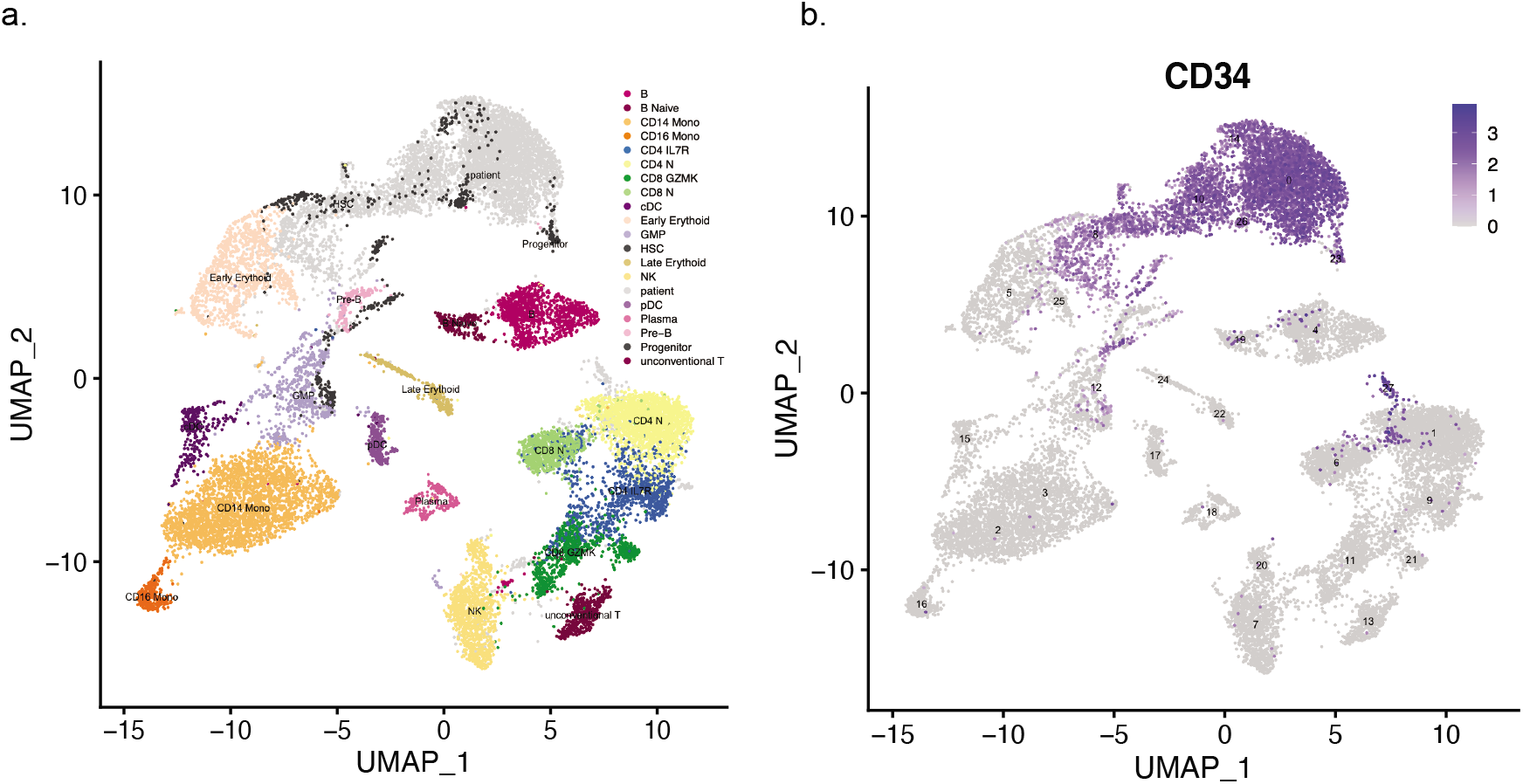
(a) Representative UMAP of computationally combining patient BM cells with healthy donor BM cells which distinguishes TME from AML cells (gray). (b) CD34 positivity labels AML cells

**Supplementary Fig. 4.**
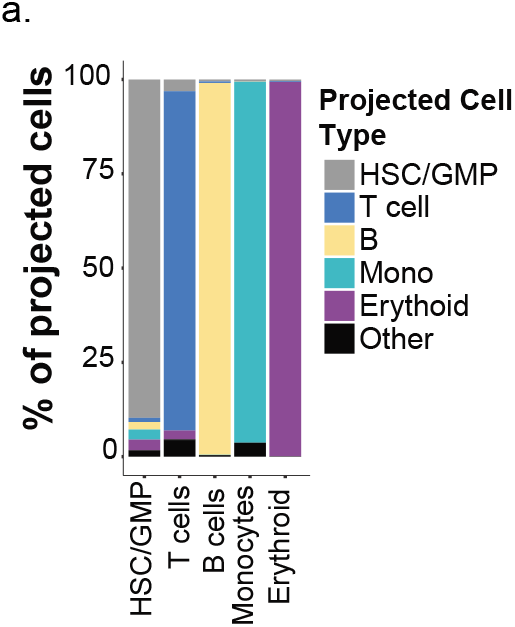
(a) Projection of TME cells onto healthy donor cells demonstrates reliable approach to identify cell lineages.

**Supplementary Fig. 5.**
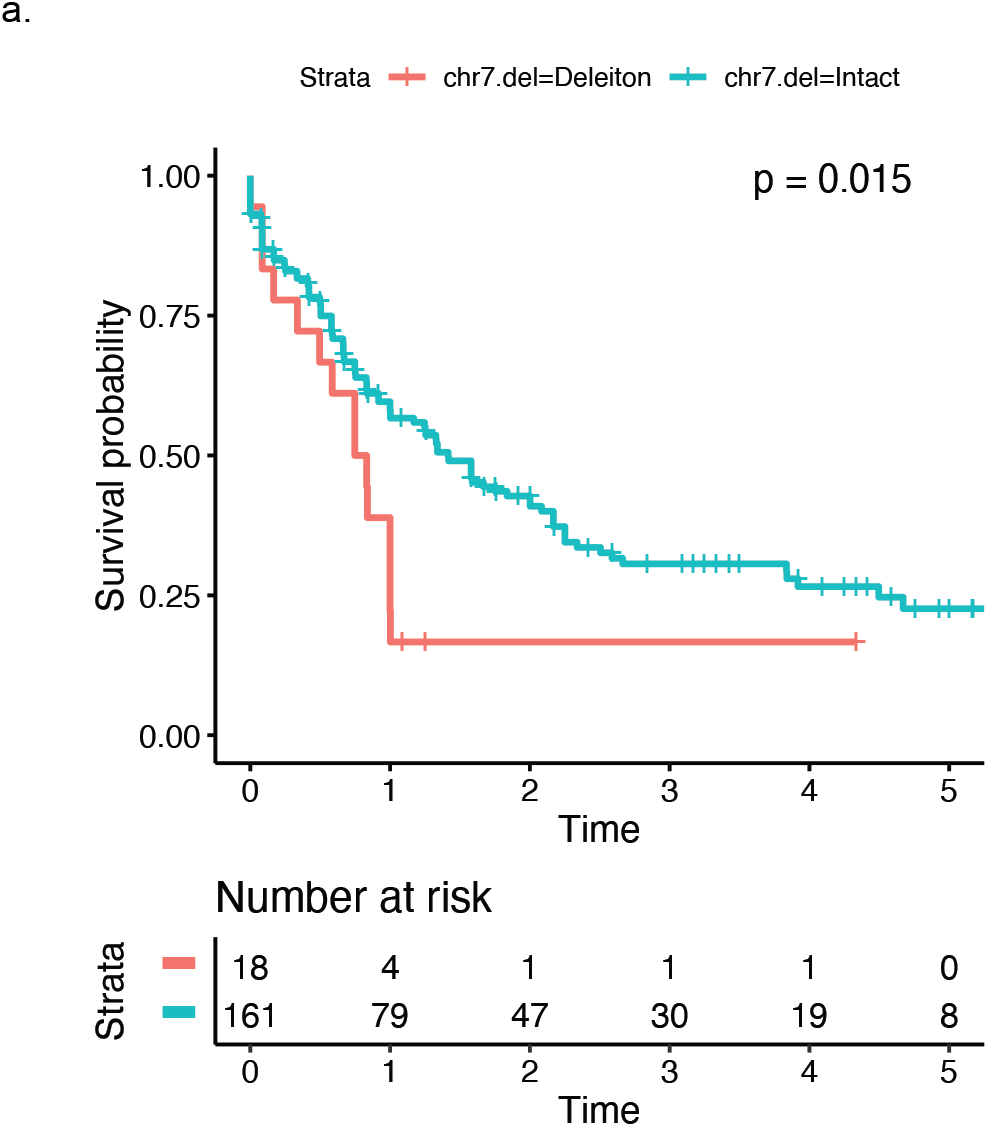
(a) Survival data of AML patients from TCGA data by chromosome 7/7q loss status.

**Supplementary Fig. 6.**
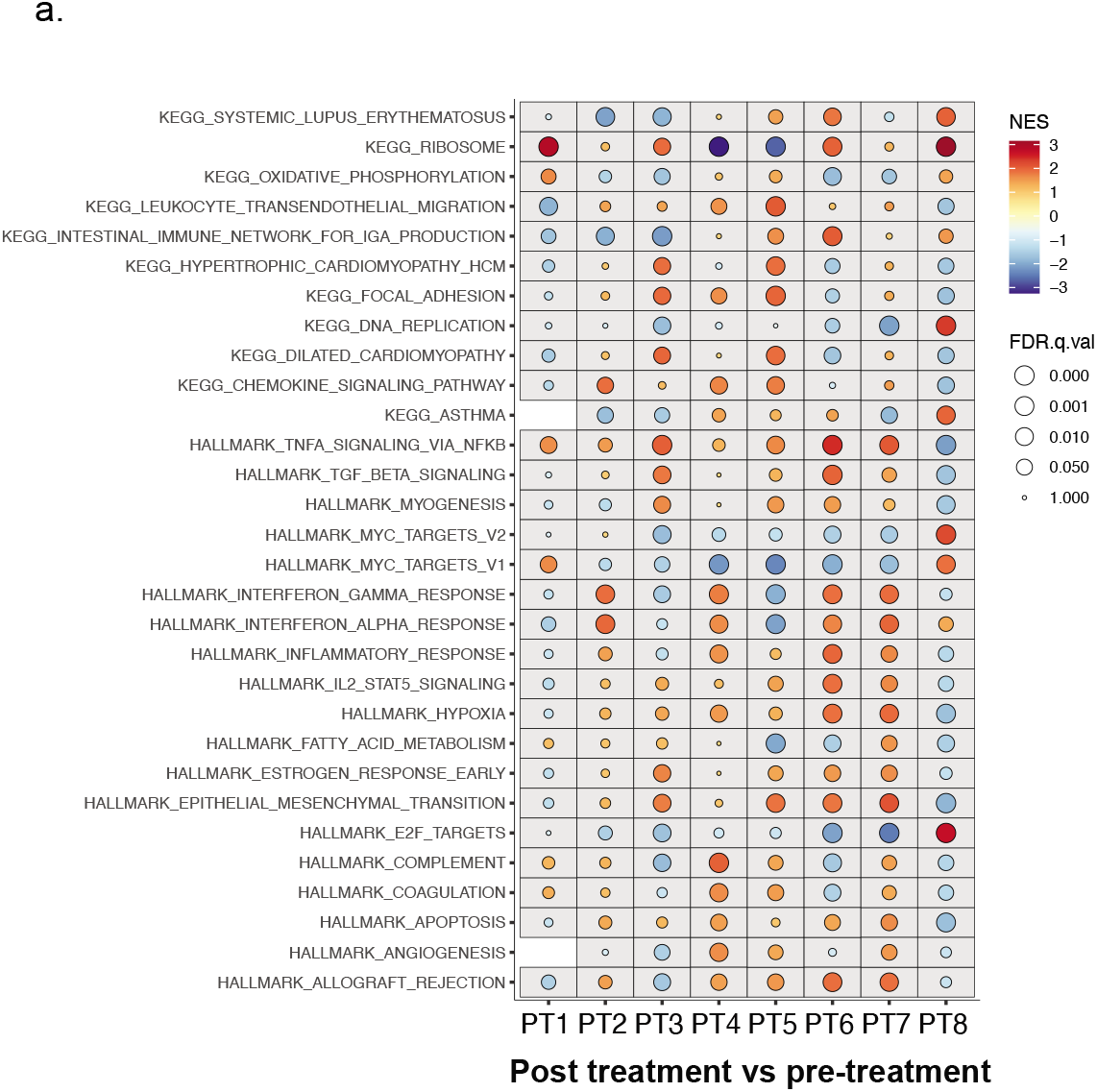
(a) Pathway enrichment of post-versus pre-treatment differentially expressed genes in AML cells demonstrating no discernible pattern.

**Supplementary Fig. 7.**
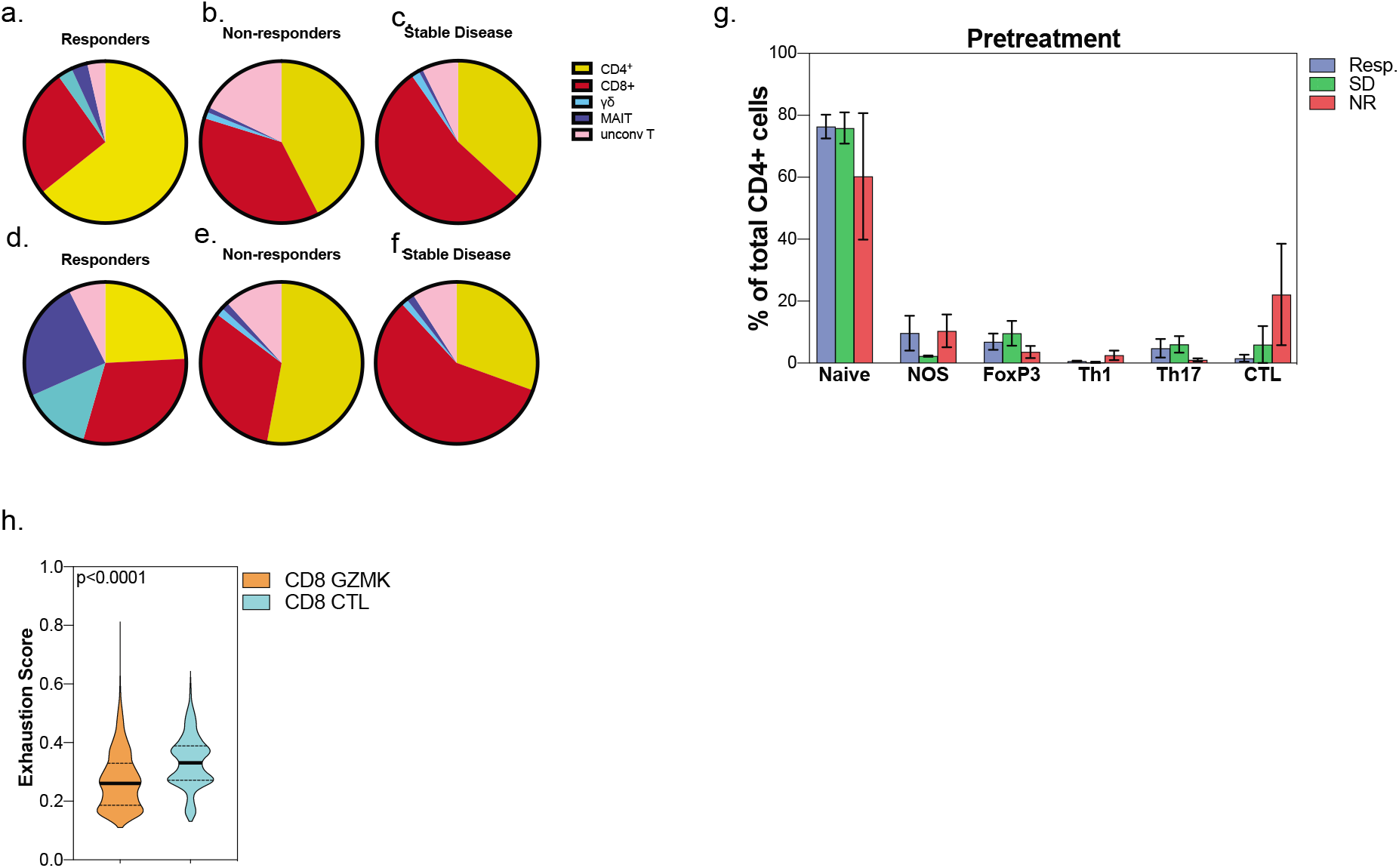
(a-f) Distribution of T cell subsets in pre and post treatment timepoints. (g) Distribution of CD4 subsets in the response groups at pre-treatment. (h) Exhaustion score of CD8 cells based on GZMK and CTL subsets.

